# Effects of the selective dopamine D_3_ receptor antagonist PG01037 on morphine-induced hyperactivity and antinociception in mice

**DOI:** 10.1101/2020.04.07.029918

**Authors:** Christian A. Botz-Zapp, Stephanie L. Foster, Desta M. Pulley, Briana Hempel, Guo-Hua Bi, Zheng-Xiong Xi, Amy Hauck Newman, David Weinshenker, Daniel F. Manvich

## Abstract

Recent preclinical studies have reported that pretreatment with the novel and highly-selective dopamine D_3_ receptor (D_3_R) antagonists R-VK4-40 or VK4-116 attenuates the abuse-related behavioral effects of oxycodone while enhancing its analgesic properties. However, whether these observed effects are generalizable to the broad class of D_3_R antagonists and/or extend to opioids other than oxycodone has not been extensively explored. The present study sought to assess the impact of pretreatment with another selective D_3_R antagonist, PG01037, on several behavioral effects of morphine in mice. C57Bl/6J mice were pretreated with PG01037 (0 – 10 mg/kg) and tested for 1) hyperlocomotion induced by acute morphine (5.6 – 56 mg/kg), 2) locomotor sensitization following repeated morphine (56 mg/kg), 3) antinociception following acute morphine (18 mg/kg), and 4) catalepsy following administration of PG01037 alone or in combination with morphine (56 mg/kg). PG01037 dose-dependently attenuated morphine-induced hyperlocomotion and morphine-induced antinociception at doses that did not alter basal locomotion or nociception alone, but did not prevent the induction of locomotor sensitization following repeated morphine administration. Moreover, PG01037 did not induce catalepsy either alone or in combination with morphine. These results suggest that attenuation of acute opioid-induced hyperlocomotion may be a behavioral effect shared among D_3_R-selective antagonists, thus supporting continued investigations into their use as potential treatments for opioid use disorder. However, PG01037 is unlike newer, highly-selective D_3_R antagonists in its capacity to reduce opioid-induced antinociception, indicating that modulation of opioid analgesia may vary across different D_3_R antagonists.

## 1 INTRODUCTION

The abuse of prescription and illicit opioids has culminated in a national healthcare crisis [1], prompting the search for novel pharmacotherapeutics that can safely and more effectively treat opioid use disorder (OUD) as compared to currently-available medications [2, 3]. The abuse-related effects of opioids are predominantly attributed to increased dopamine (DA) neurotransmission within the mesolimbic reward system [for review, see 4, 5, 6], a projection arising from DAergic neurons located in the ventral tegmental area (VTA) and terminating in the nucleus accumbens (NAc) [7, 8]. Opioids administered either systemically [9, 10] or directly into the VTA [10–13] produce increases in NAc DA levels by disinhibiting VTA DA neurons [14, 15]. Accordingly, the locomotor-activating, reinforcing, and reinstatement-inducing effects of opioids are each dampened following perturbation of NAc DA neurotransmission [16–23]. DA binds to five G protein-coupled receptor subtypes which are divided into two families. The D_1_-like receptor family includes the G_s_-coupled D_1_ and D_5_ receptor subtypes (D_1_R and D_5_R) while the D_2_-like receptor family includes the G_i_-coupled D_2_, D_3_, and D_4_ receptor subtypes (D_2_R, D_3_R, D_4_R) [24]. Administration of nonselective antagonists at either D_1_-like receptors or D_2_-like receptors reduces opioid-induced locomotor activation, opioid self-administration, and opioid seeking [for review, see 4, 5, 6, 25]. However, adverse side effects have hampered the potential clinical utility of these drug classes as treatments for substance use disorders [26–29]. Attention has therefore shifted towards receptor subtype-selective compounds that may retain pharmacotherapeutic efficacy for the treatment of OUD while lacking undesirable behavioral effects.

The D_3_R has emerged as an appealing pharmacological target in this regard as reviewed elsewhere [30–32]. Of most relevance to this report is preclinical evidence that selective D_3_R antagonism attenuates opioid-induced hyperactivity, opioid self-administration, and opioid-seeking behavior, without producing adverse motoric effects associated with nonselective D_2_-like receptor antagonism [33–39]. Furthermore, two newly-developed and highly-selective D_3_R antagonists, R-VK4-40 and VK4-116, were recently reported to enhance the analgesic effects of oxycodone while simultaneously reducing its abuse-related effects, lending additional support to the potential use of D_3_R antagonists as treatments for OUD and also possibly as add-on medications to be administered concurrently with opioid analgesics [37, 38]. However, no other studies to date have investigated the impact of highly-selective D_3_R antagonists on opioid-induced analgesia [32], leaving unresolved whether the analgesia-enhancing effect observed with R-VK4-40 and VK4-116 extends to other D_3_R antagonists and/or to analgesia induced by opioids other than oxycodone.

To begin to address these questions, the present study sought to determine the effects of pretreatment with another selective D_3_R antagonist, PG01037 (133-fold selectivity for D_3_R over D_2_R [40]), on various unconditioned behavioral effects of morphine. PG01037 was selected for use in these studies for two major reasons. First, the impact of PG01037 administration on any opioid-mediated effect has not previously been investigated [32]. Second, existing evidence already suggests that PG01037 may differentially modulate the behavioral effects of drugs of abuse as compared to other selective D_3_R antagonists. For example, PG01037 pretreatment significantly enhances cocaine-induced hyperlocomotion [41], whereas other highly-selective D_3_R antagonists either attenuate or have no effect on this behavioral response [42]. We therefore reasoned that PG01037 would be an ideal test compound with which to assess whether modulations of the behavioral effects of opioids might also be dissimilar among D_3_R-selective antagonists.

Stimulation of locomotor activity in rodents is a useful and straightforward unconditioned behavioral response with which to interrogate abuse-related increases in NAc DA neurotransmission following systemic administration of drugs of abuse, including opioids [43–47]. Previous work has demonstrated that pretreatment with the highly-selective D_3_R antagonists YQA14 or VK4-116 attenuates morphine- and oxycodone-induced hyperlocomotion respectively in mice [36, 48], but whether PG01037 similarly disrupts opioid-induced locomotor increases has not been investigated. We therefore first assessed the impact of PG01037 pretreatment on acute morphine-induced hyperactivity as well as the induction of locomotor sensitization to repeated morphine administration in mice. We next examined whether PG01037 modulates the antinociceptive effects of morphine. Finally, because nonselective blockade of D_2_-like receptors produces catalepsy alone and potentiates opioid-induced catalepsy [49–51], we investigated whether administration of PG01037 alone, or in combination with morphine, would induce cataleptic effects.

## 2 MATERIALS AND METHODS

### 2. 1 Subjects

Subjects used in this study were 64 adult male and female C57BL/6J mice (32/sex), 8-12 weeks old at the start of study. Mice were either acquired from Jackson Laboratory (Bar Harbor, ME; n = 40) or from a breeding colony at the National Institute on Drug Abuse (n = 24). Mice were housed in same-sex groups of 3-5 per cage in a climate-controlled vivarium with a 12-hr light cycle and had *ad libitum* access to food and water in the home cage. Procedures were conducted in accordance with the *Guide for the Care and Use of Laboratory Animals* of the U.S. National Research Council and were approved by Institutional Animal Care and Use Committees at Emory University, Rowan University, or the National Institute on Drug Abuse of the National Institutes of Health. All behavioral testing was performed during the light cycle.

### 2.2 Locomotor Activity Apparatus

Locomotor activity was assessed in transparent polycarbonate cages (22 × 43 × 22 cm) that allowed passage of 8 infrared beams through the long wall and 4 infrared beams through the short wall of the enclosure at 4.9 cm intervals (San Diego Instruments; San Diego, California). Horizontal ambulations, defined as the sequential disruption of two adjacent infrared beams, were recorded in 5-min bins. The test chambers were prepared with a thin layer of clean bedding prior to each test session. Before the onset of experiments, mice were injected i.p. with saline and placed in the test chambers for 30 min for 3 consecutive days in order to habituate the mice to injections and the test apparatus.

### 2.3 Acute Morphine-Induced Locomotion

The effects of PG01037 on acute morphine-induced locomotion were evaluated in 16 mice (8 males, 8 females) using a within-subjects design. Animals were initially placed in the center of the locomotor chamber and ambulations were recorded for 90 min. Next, animals were briefly removed from the chamber, injected with PG01037 (vehicle, 0.1, 1, or 10 mg/kg i.p.), and returned to the locomotor chamber for 30 min. Finally, mice were again removed from the chamber, injected with morphine (vehicle, 5.6, 18, or 56 mg/kg i.p.), and placed back in the chamber for 120 min. The dose range for PG01037 was carefully selected for use based on our previous work showing that doses up to 10 mg/kg do not appreciably disrupt basal locomotion but significantly modulate the locomotor-activating effects of cocaine in C57BL/6J mice [41]. The 5.6 – 56 mg/kg dose range of morphine was selected based on our own pilot studies showing that it captures both the ascending and descending limbs of the morphine dose-response curve. Dose administration of PG03017 was pseudorandomized and counterbalanced across animals within each dose of morphine. All doses of PG01037 were assessed for a given morphine dose before switching to a different morphine dose. The order of morphine dose testing was 18 mg/kg, 56 mg/kg, 5.6 mg/kg, vehicle. All test sessions were separated by at least 1 week to prevent the development of locomotor sensitization to morphine. All mice received all treatments.

### 2.4 Morphine-Induced Locomotor Sensitization

Sensitization induction took place over 5 consecutive days and was performed in 2 separate groups of mice (n = 8/group, 4 male and 4 female). Mice were initially placed in the center of the locomotor chamber, and locomotor activity was recorded for 90 min. They were then briefly removed from the chamber, injected with PG01037 (vehicle or 10 mg/kg, i.p.), and returned to the locomotor chamber for 30 min. Mice were again removed and injected with morphine (56 mg/kg i.p.), then placed back in the chamber for 120 min. Mice received the same dose of PG01037 across each of the 5 induction days. Seven days following the last induction session, locomotor activity was again assessed as described above with the exception that all mice received vehicle as the pretreatment 30 min prior to challenge with 56 mg/kg morphine.

### 2.5 Hot Plate Test for Thermal Nociception

Antinociception was assessed in mice using a hot plate system (Model 39, IITC Life Science Inc., Woodland Hills, CA, USA) set to 52 ± 0.2°C. Mice were placed on the platform surrounded by transparent Plexiglas walls and removed after the first sign of thermal distress (paw licking, jumping, hind paw stomping). The latency to the first indicator of pain was recorded. A maximal cutoff of 60 s was instituted to prevent tissue damage. The antinociceptive effects of PG01037 alone were assessed in one group of 8 mice (4 males, 4 females). Subjects were first placed on the hot plate prior to any drug treatment to measure baseline response latencies (time point 0). Next, mice were administered PG01037 (vehicle, 0.1, 1, or 10 mg/kg, i.p.) and tested on the hot plate at 30, 60, 90, and 120 min post-injection. PG01037 doses were counterbalanced across subjects. The effects of PG01037 pretreatment on morphine-induced antinociception were examined in a separate group of 16 mice (8 males, 8 females). Following baseline testing (time point 0), mice were administered PG01037 (vehicle, 0.1, 1, or 10 mg/kg, i.p.) followed 30 min later by morphine (18 mg/kg i.p.). Hot plate testing was assessed at 30, 60, 90, and 120 min post-morphine injection. Each mouse received 1-3 doses of PG01037 in a counterbalanced manner whereby all possible PG01037 × morphine dose combinations consisted of n = 8. For each experiment, hot plate test sessions were separated by 2-3 days.

### 2.6 Catalepsy

The capacity of PG01037 alone or in combination with morphine to produce catalepsy was assessed in 8 mice (4 males, 4 females). Mice received a pretreatment of either PG01037 (vehicle or 10 mg/kg, i.p.) followed 30 min later by morphine (vehicle or 56 mg/kg, i.p.). Catalepsy was evaluated using the “bar test” [52], during which a thin bar was secured horizontally 1.75 in above a flat surface. Each test was conducted by lifting the mouse by the tail and allowing it to grab the bar with its front paws, then releasing the tail so that the mouse was positioned sitting upright on its hind legs. Upon assuming this position, the latency to remove at least one paw from the bar was recorded. The test was stopped if the subject failed to withdraw one paw within 60 s. Mice that could not be placed in the testing position after 3 attempts received a latency score of 0 s. In each test, catalepsy was measured 0, 15, 30, 60, and 120 min following administration of morphine. The order of dose-combinations was randomized across mice. Mice were tested once per week until each mouse received all treatment combinations. After PG01037/morphine testing was completed, all mice received a final catalepsy test in which they were administered risperidone (3 mg/kg, i.p.) followed by saline i.p. 30 min later. Catalepsy in these tests was measured up to 60 min following saline injection. The 3 mg/kg risperidone test was included as a positive control as it induces prominent catalepsy in mice [53].

### 2.7 Drugs

Morphine sulfate (National Institute on Drug Abuse Drug Supply Program, Bethesda, MD) was dissolved in sterile saline. PG01037 was synthesized by Ms. J. Cao in the Medicinal Chemistry Section, National Institute on Drug Abuse Intramural Research Program as described previously [40] and dissolved in sterile water. Risperidone (Sigma-Aldrich; St. Louis, MO) was dissolved in vehicle containing ethanol:CremophorEL (Sigma-Aldrich):saline (5:10:85 v/v). All drugs were administered i.p. at a volume of 10 ml/kg.

### 2.8 Statistical Analyses

For acute morphine-induced locomotion studies, total ambulations during the 2 h following morphine administration were analyzed via two-way ANOVA with repeated measures on both factors (PG01037 dose × morphine dose), followed by post hoc Dunnett’s multiple comparisons tests to compare each dose of PG01037 to its vehicle within each dose of morphine. Locomotor activity in the 30-min period after PG01037 administration (prior to morphine administration) was analyzed using one-way repeated measures ANOVA. Effects of vehicle or 10.0 mg/kg PG01037 alone on locomotor activity were assessed by paired t-test. For the induction phase of sensitization (i.e. days 1-5), total ambulations during the 2 h following morphine administration were analyzed via mixed two-way ANOVA with repeated measures on one factor (day) and independent measures on the other factor (PG01037 dose). Dunnett’s multiple comparisons tests were used to compare morphine-induced locomotor activity on each of induction days 2-5 vs. day 1. For the challenge day of sensitization studies (day 12), total ambulations during the 2 h following morphine administration were analyzed via independent two-tailed t-test. The antinociceptive effects of PG01037 alone or in combination with morphine were analyzed using a two-way ANOVA with repeated measures on one factor (time) and independent measures on the other factor (PG01037 dose), followed by Dunnett’s or Tukey’s multiple comparisons tests, as specified in the text. Latency scores in catalepsy experiments were analyzed using two-way ANOVA with repeated measures on both factors (treatment × time). The effect of 3 mg/kg risperidone + saline was excluded from statistical analyses because risperidone was included only as a positive control to validate the catalepsy detection procedure. All data were plotted and analyzed using GraphPad Prism v8.4 (GraphPad Software, La Jolla, CA, USA). Significance was set at *p* < 0.05 for all tests.

## 3 RESULTS

### 3.1 Effects of PG01037 on Acute Morphine-Induced Hyperlocomotion

Administration of morphine after vehicle pretreatment resulted in increased locomotor activity with a typical inverted U-shaped dose-response function (Fig. 1). Two-way repeated measures ANOVA of PG01037 in combination with 5.6 – 56 mg/kg morphine indicated significant main effects of morphine dose (F_(2,30)_ = 5.82, p = 0.007), PG01037 dose (F_(3,45)_ = 29.27, p < 0.0001), and a significant morphine × PG01037 interaction (F_(6,90)_ = 5.90, p < 0.0001). Post hoc comparisons revealed that pretreatment with 0.1 mg/kg PG01037 slightly elevated hyperlocomotion produced by 18 mg/kg morphine, whereas higher doses of PG01037 dose-dependently and more robustly attenuated the effects of 18 and 56 mg/kg morphine. The inhibitory actions of PG01037 on morphine-induced locomotion typically emerged within 5-15 min following morphine administration and persisted for the duration of the 120-min observation period (Fig. 2A-C). PG01037 administration did not significantly alter locomotor activity in the 30 min following its administration, prior to morphine injection (one-way repeated measures ANOVA, (F_(3,45)_ = 0.33, p = 0.801) (Fig. S1). To further confirm a lack of effect by PG01037 alone, vehicle or 10 mg/kg PG01037 were administered followed 30 min later by saline and locomotion was monitored for 120 min. 10 mg/kg PG01037 pretreatment did not alter total ambulations during this longer observation period (paired t-test, t_(15)_ = 1.58, p = 0.13) (Fig. S2A-B).

**Fig. 1.**
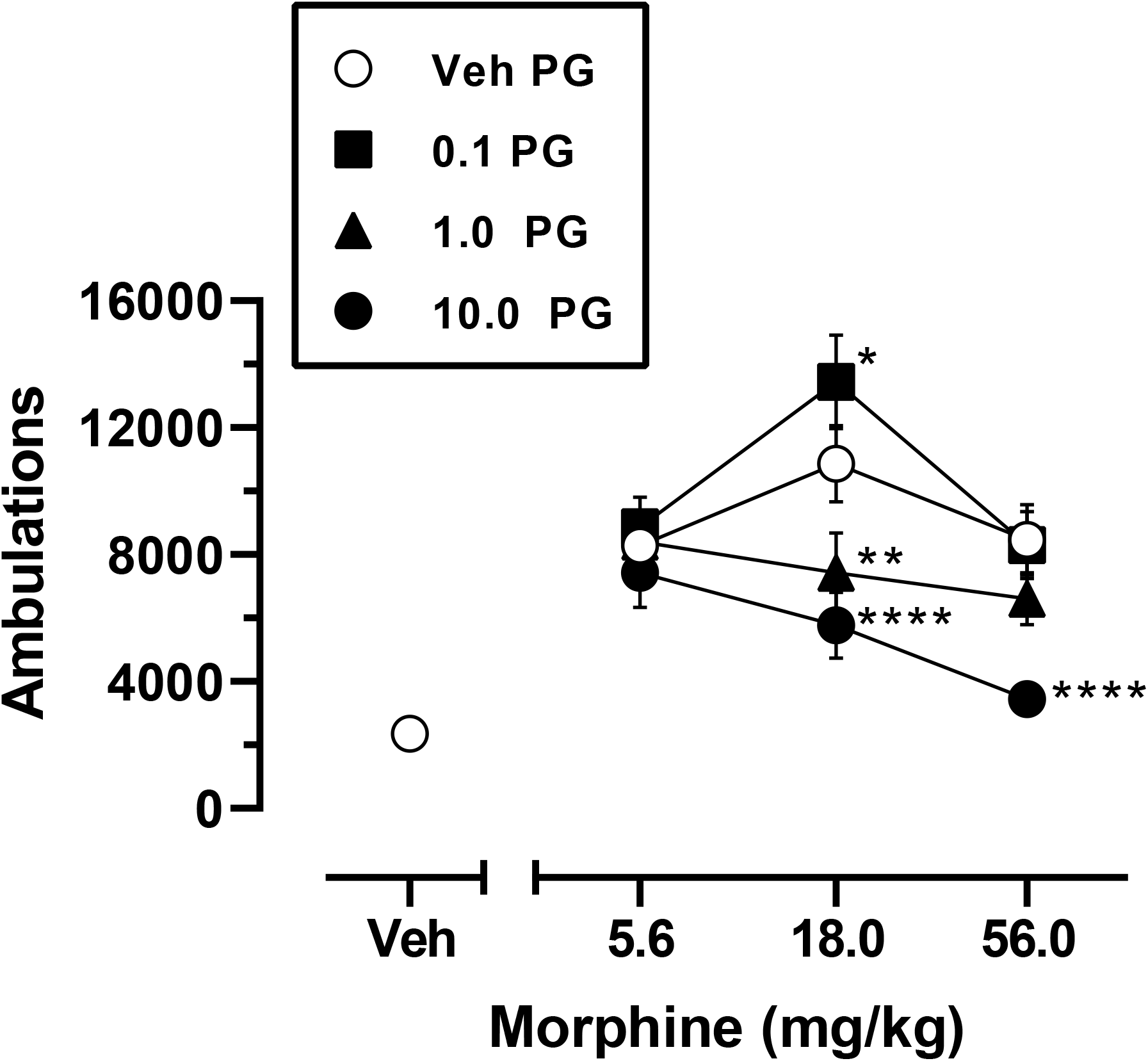
Effects of pretreatment with PG01037 on acute morphine-induced locomotor activity. Mice were pretreated with vehicle or 0.1 – 10 mg/kg PG01037, followed 30 min later by vehicle or 5.6 – 56 mg/kg morphine. Shown are mean ± SEM total ambulations in the 120-min period following morphine administration. * p < 0.05, ** p < 0.01, **** p < 0.0001, compared to vehicle at the same dose of morphine. All mice received all treatments (n = 16). “Veh” = vehicle; “PG” = PG01037. Doses on the abscissa are plotted along a log scale.

**Fig. 2.**
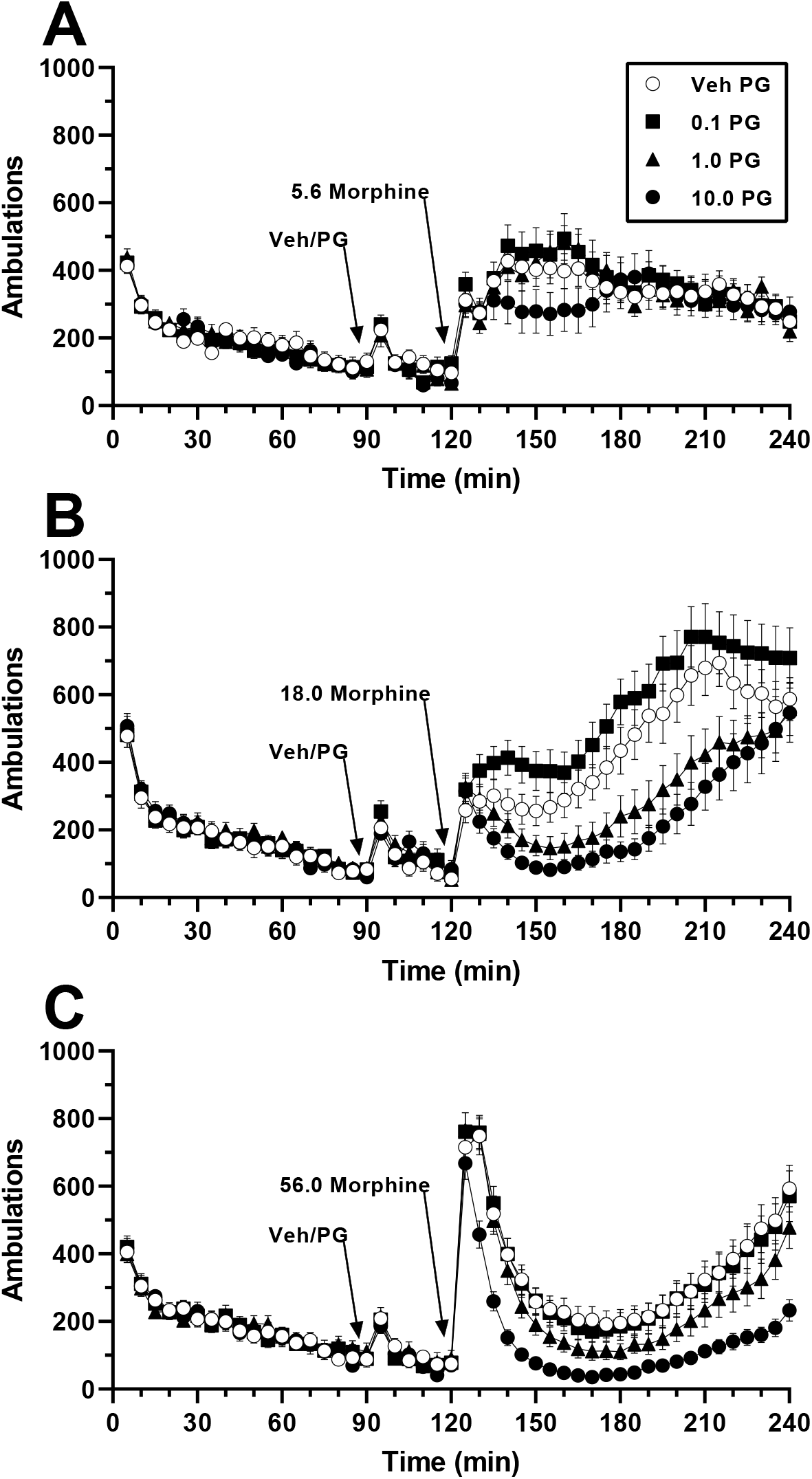
Time course of changes in locomotor activity following pretreatment with PG01037 and subsequent administration of morphine. PG01037 (vehicle, 0.1 – 10 mg/kg) was administered 30 min prior to **A** 5.6 mg/kg morphine, **B** 18 mg/kg morphine, or **C** 56 mg/kg morphine. Each data point represents mean ± SEM ambulations recorded in 5-min bins. Arrows indicate time of pretreatment injection (“Veh/PG”, i.e. vehicle or 0.1 – 10 mg/kg PG01037 respectively) or time of morphine injection. All mice received all treatments (n = 16). “Veh” = vehicle; “PG” = PG01037.

### 3.2 Effects of PG01037 on Morphine-Induced Locomotor Sensitization

To test the impact of selective D_3_R antagonism on the development of morphine-induced locomotor sensitization, mice were pretreated daily for 5 consecutive days with vehicle or PG01037 (10 mg/kg i.p.) 30 min prior to morphine (56 mg/kg, i.p.). 10 mg/kg PG01037 was selected for use in this experiment because it produced the greatest attenuation of morphine’s acute locomotor activity and did not disrupt basal locomotion in the preceding experiment. Two-way mixed factors ANOVA revealed significant main effects of induction day (F_(4, 56)_ = 17.66, p < 0.0001) and PG01037 dose (F_(1, 14)_ = 4.67, p = 0.049), but not a significant day × PG01037 interaction (F_(4, 56)_ = 0.51, p = 0.73). Post hoc analyses indicated that, collapsed across vehicle and 10 mg/kg PG01037 pretreatments, mice exhibited a sensitized locomotor response to morphine by day 3 of the induction phase (Fig. 3; time course, Fig. 4A-E). The significant main effect of PG01037 treatment without a significant day × PG01037 interaction indicates that PG01037 attenuated morphine-induced hyperlocomotion equally across all 5 days, effectively reducing it on average by ~ 36.5% (Fig. 3; time course, Fig. 4A-E). One week after the final induction session, all mice received vehicle followed by a morphine challenge (56 mg/kg, i.p.). Independent t-test showed no difference in locomotor activity between mice that had previously been pretreated with 10 mg/kg PG01037 during induction as compared to mice pretreated with vehicle during induction (t(14) = 0.67, p = 0.52) (Fig. 3; time course, Fig. 4F).

**Fig. 3.**
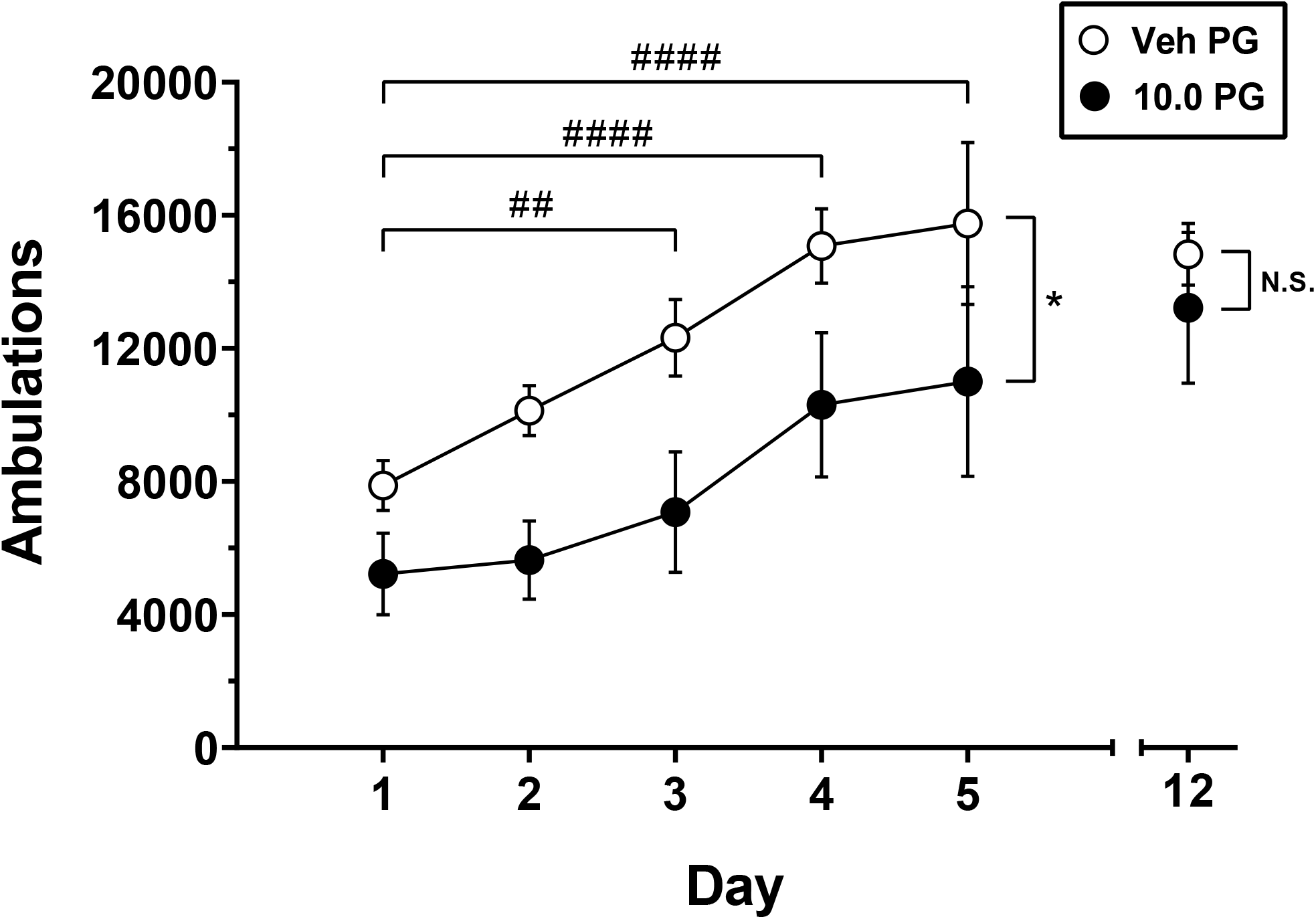
Effects of pretreatment with PG01037 on morphine-induced locomotor sensitization. Mice received the combination of either vehicle + 56 mg/kg morphine or 10 mg/kg PG01037 + 56 mg/kg morphine daily for 5 days. One week later (day 12), all mice received vehicle pretreatment prior to a challenge with 56 mg/kg morphine. Shown are mean ± SEM total number of ambulations in the 120-min period following injection of 56 mg/kg morphine, which was administered 30 min after PG01037 pretreatment. ## p < 0.01, #### p < 0.0001, significant difference compared to Day 1 (collapsed across PG01037 pretreatment doses). * p < 0.05, main effect of pretreatment dose (collapsed across days 1-5). n = 8/group. “N.S.” = not significant; “Veh” = vehicle; “PG” = PG01037.

**Fig. 4.**
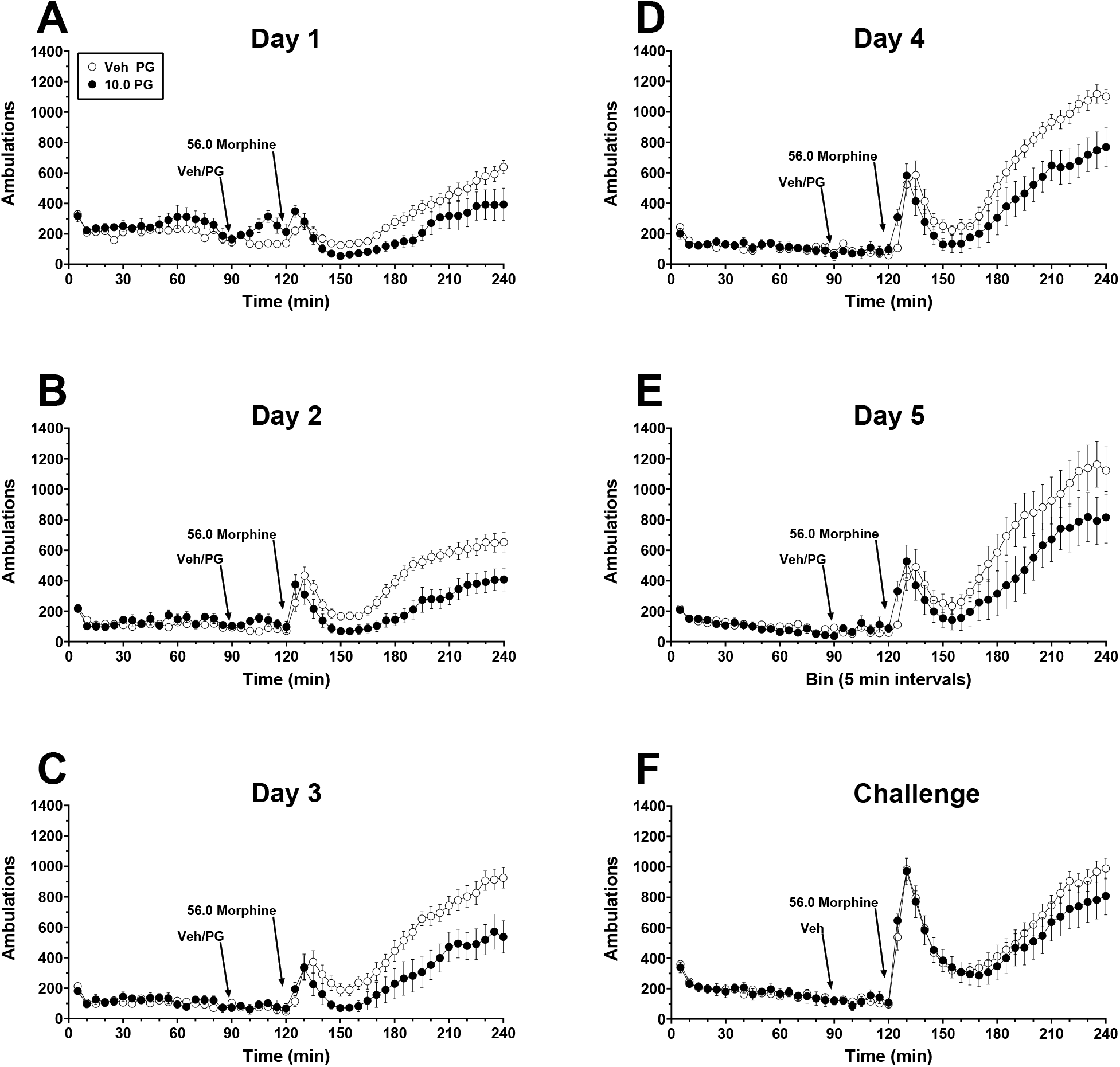
Time course of locomotor activity following pretreatment with PG01037 and subsequent administration of 56 mg/kg morphine during induction days 1-5 of sensitization, and challenge test with morphine alone one week later. Experimental details are as described for Figure 3. Shown are mean ± SEM ambulations recorded in 5-min bins on **A** induction day 1, **B** induction day 2, **C** induction day 3, **D** induction day 4, and **E** induction day 5 of sensitization induction, or **F** challenge day. Arrows indicate time of pretreatment injection (“Veh/PG”, i.e. vehicle or 10 mg/kg PG01037 respectively) or time of morphine injection. n = 8/group. “Veh” = vehicle; “PG” = PG01037.

### 3.3 Effects of PG01037 on Morphine-Induced Antinociception

The effects of PG01037 administration alone on nociception in the hot plate test are shown in Fig. 5A. Two-way mixed-factors ANOVA indicated a significant main effect of time (F_(4, 112)_ = 4.02, p = 0.004) but not of PG01037 dose (F_(3, 28)_ = 0.82, p = 0.49) or a time × PG01037 interaction (F_(12, 112)_ = 0.18, p = 0.18). Post hoc Tukey’s tests revealed that, collapsed across PG01037 dosing conditions, reaction latencies decreased slightly but significantly at the 30-min and 60-min time points as compared to 0 min (p < 0.05).

**Fig. 5.**
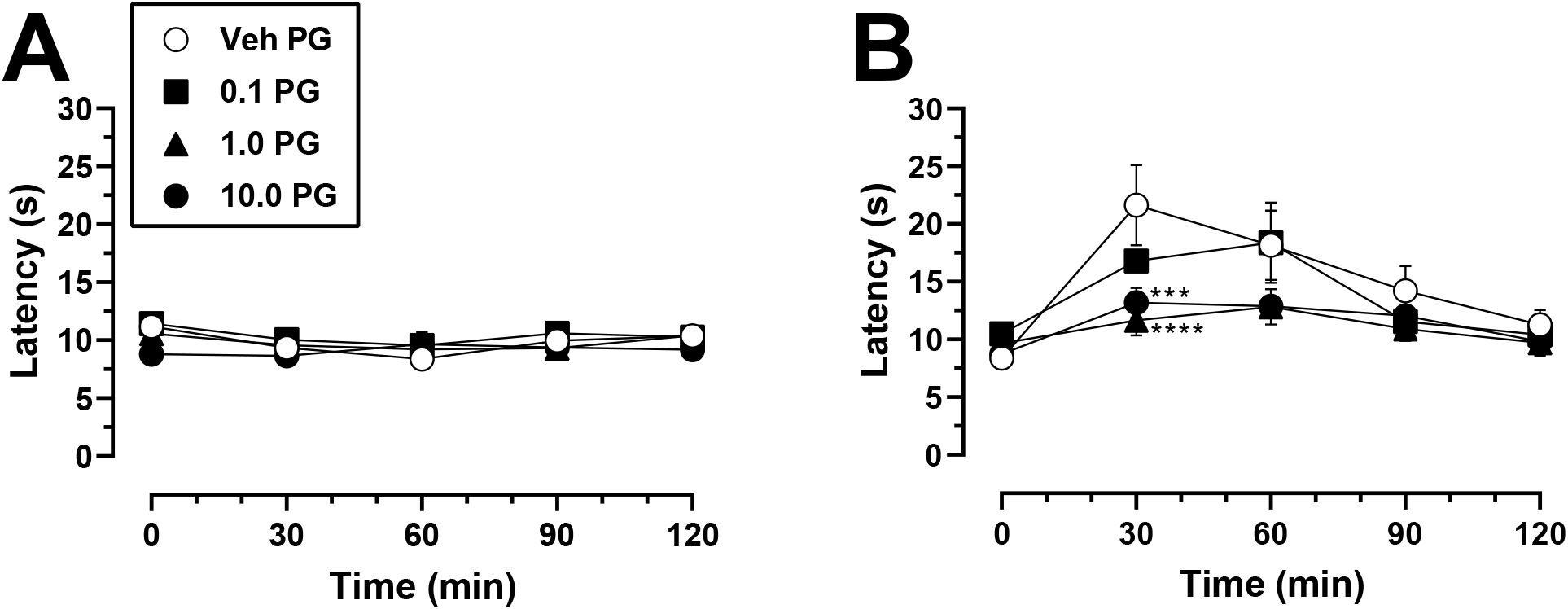
Thermal nociception in mice following pretreatment with PG01037 alone or in combination with morphine. **A** Mice were administered PG01037 (vehicle, 0.1 – 10 mg/kg) and thermal nociception was assessed over 120 min following PG01037 injection. **B** Mice were administered PG01037 (vehicle, 0.1 – 10 mg/kg) 30 min prior to 18 mg/kg morphine, and thermal nociception was assessed over 120 min following morphine injection. Each data point represents mean ± SEM latency in seconds to the first indicator of nociception. *** p < 0.001, **** p < 0.0001, compared to vehicle at the same dose of morphine. n = 8/group. “Veh” = vehicle; “PG” = PG01037.

In mice pretreated with vehicle, 18 mg/kg morphine increased reaction latency with a maximal efficacy of ~2.5-fold evident 30 min after morphine administration that gradually returned to near-baseline levels by the 120-min time point (Fig. 5B). Two-way mixed factors ANOVA indicated a significant main effect of time (F_(4, 112)_ = 23.41, p < 0.0001), no main effect of PG01037 dose (F_(3, 28)_ = 2.29, p = 0.10), and a significant time × PG01037 interaction (F_(12, 112)_ = 2.77, p = 0.003). Post hoc Dunnett’s tests revealed that compared to vehicle pretreatment, administration of 1 or 10 mg/kg PG01037 significantly attenuated the antinociceptive effects of morphine at the 30-min time point when morphine’s effects were maximal, reducing them by 40% and 54%, respectively. This attenuating effect of PG01037 fell just short of statistical significance at the 60-min time point (p = 0.06 for both 1 and 10 mg/kg PG01037 compared to vehicle).

### 3.4 Effects of PG01037 Alone or in Combination with Morphine on Catalepsy

To determine whether selective D_3_R antagonism induces catalepsy either alone or in combination with morphine, mice were administered PG01037 (vehicle or 10 mg/kg, i.p.) 30 min prior to morphine (vehicle or 56 mg/kg, i.p.). We purposely selected the highest doses administered of each compound in our locomotor and nociception experiments in order to maximize the potential detection of catalepsy. Neither administration of PG01037 alone, morphine alone, nor their combination resulted in catalepsy (Fig. 6). Analysis of these treatment conditions using two-way repeated measures ANOVA showed no significant main effect of treatment (F_(2,14)_ = 1.00, p = 0.39), time (F_(4,28)_ = 1.00, p = 0.42), or a treatment × time interaction (F_(8,56)_ = 1.00, p = 0.45). For all mice tested, the latency to withdraw a forepaw in any of the aforementioned conditions did not exceed 1 s. By contrast, 3 mg/kg risperidone produced a robust increase in catalepsy across the 60-min test period, ranging on average from ~ 48 s up to the procedural maximum time allowed of 60 s (Fig. 6).

**Fig. 6.**
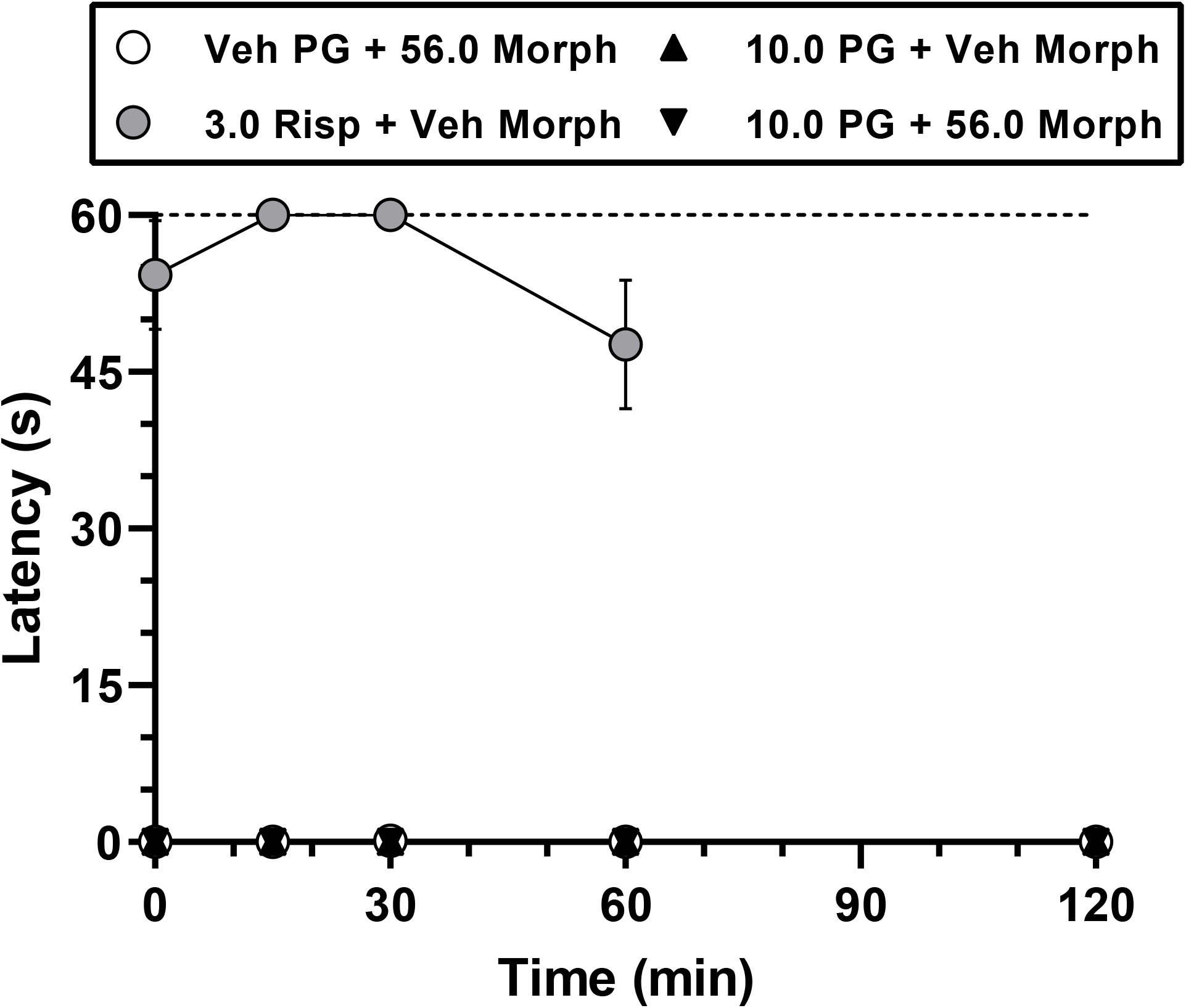
Catalepsy following administration of PG01037 alone or in combination with morphine. Mice were pretreated with PG01037 (vehicle or 10 mg/kg) followed 30 min later by administration of vehicle or 56 mg/kg morphine, or 3 mg/kg risperidone followed 30 min later by administration of vehicle morphine. Each data point represents mean ± SEM latency in seconds to withdraw a paw in the bar test. Latencies were measured at 0, 15, 30, 60, and 120 min relative to the second injection. The dotted line represents the 60-s maximal time allowed for paw withdrawal. All mice received all treatments (n = 8). “Veh” = vehicle; “PG” = PG01037; “Risp” = risperidone; “Morph” = morphine.

## 4 DISCUSSION

In the present study, pretreatment with PG01037 dose-dependently attenuated acute morphine-induced hyperactivity in mice. This finding is in agreement with previous reports demonstrating that pretreatment with several other selective D_3_R antagonists similarly produces significant reductions in the locomotor-activating effects of morphine or oxycodone [36, 48, 54]. Collectively, these results suggest that D_3_R antagonism reliably attenuates the locomotor-activating effects of opioids regardless of the specific compound used, indicating a D_3_R antagonist class effect. Because the locomotor-activating effects of opioids are most often attributed to increased DA neurotransmission within the mesolimbic system [17, 43, 45], the attenuated locomotor response to opioids that is produced by D_3_R antagonists may reflect as-yet unidentified modulations in mesolimbic DA signaling and/or NAc output that are likely to also mediate their concomitant reductions in opioid reward. It is interesting to note that while various D_3_R antagonists all appear to attenuate opioid-induced hyperlocomotion, their impact on psychostimulant-induced hyperlocomotion is more variable. We and others have reported that PG01037 and the selective D_3_R antagonist NGB294 enhance the locomotor-activating effects of cocaine or amphetamine respectively [41, 55], whereas other D_3_R antagonists either reduce or do not affect psychostimulant-induced hyperlocomotion [42, 56, 57]. The reasons as to why D_3_R antagonists reliably attenuate opioid-induced hyperlocomotion but exhibit more diverse effects on stimulant-induced hyperlocomotion remain unclear and will require further research to resolve.

In contrast to its effects on acute morphine-induced hyperactivity, PG01037 did not disrupt the induction of locomotor sensitization in the present study, since PG01037-treated mice displayed sensitized responses to morphine that were comparable to vehicle-treated mice. Although the NAc contributes to the locomotor-activating effects of acute systemic opioid administration [58–60], it is generally accepted that neuroadaptations within the VTA, and not the NAc, underlie the development of opioid-induced locomotor sensitization [61–63]. The finding that PG01037 pretreatment attenuates acute morphine-induced hyperactivity but not its sensitization may therefore indicate that PG01037 reduces opioid-induced hyperlocomotion via actions in the NAc that effectively “mask” the overt appearance of sensitization, whereas the VTA-dependent adaptations underlying sensitization are unaltered by PG01037 and can be “unmasked” when subjects are challenged with morphine alone. Additional studies assessing the impact of intra-VTA or intra-NAc administration of PG01037 and/or other D_3_R antagonists on the induction of opioid locomotor sensitization will be required to test the veracity of this hypothetical schema. It is noteworthy that our sensitization results with PG01037 seem to contradict a report that pretreatment with the highly-selective D_3_R antagonist VK4-116 attenuated the induction of oxycodone-induced locomotor sensitization [48]. However, some key procedural differences may underlie these discrepant findings including the use of different opioids (morphine vs. oxycodone), imposition of 7 days vs. 2 days between the final induction session and the expression test, or most notably, use of different D_3_R antagonists (PG01037 vs. VK4-116) which may themselves exert different qualitative effects on opioid-induced locomotor sensitization for reasons not yet understood.

Whereas PG01037 administration alone did not disrupt thermal nociception in the present study, it dose-dependently attenuated the antinociceptive effects of acute morphine, evidenced by an apparent downward shift of morphine’s efficacy over time. This result is in opposition to recent studies using the newer and highly-selective D_3_R antagonists VK4-116 and R-VK4-40, as pretreatment with these compounds enhances the antinociceptive effects of oxycodone in rats [37, 38]. Why PG01037 produces an opposite effect on opioid-mediated antinociception in the present study remains unclear, although VK4-40 and VK4-116 notably exhibit some other effects that are discordant with older D_3_R antagonists; for example, they do not potentiate the cardiovascular effects of cocaine as compared to older-generation D_3_R antagonists [42]. It is also noteworthy that our present study examined the antinociceptive effects of morphine rather than oxycodone, and did so in mice rather than rats. These procedural differences aside, our present findings highlight the need for more research to clarify the mechanisms by which highly-selective D_3_R antagonists modulate the analgesic properties of clinically-utilized opioid analgesics [32], and to ascertain why different antagonists may yield different results.

Combined administration of nonselective D_2_-like receptor antagonists with opioids induces catalepsy in mice that is substantially greater than that produced by either drug alone [49, 50]. It is generally believed that D_2_-like receptor antagonists exert these effects via actions at the D_2_R subtype because selective D_2_R blockade alone produces catalepsy in mice [64], rats [65], and nonhuman primates [66], whereas catalepsy has not been detected following treatment with D_3_R antagonists [65, 67]. In agreement with these latter findings, PG01037 in the present study showed no evidence of inducing cataleptic effects alone at a behaviorally-relevant dose that significantly modulated morphine-induced hyperactivity and antinociception. More importantly, the present study is the first to demonstrate that concurrent administration of high doses of a selective D_3_R antagonist (PG01037) and an opioid (morphine) does not induce catalepsy. Given that D_3_R antagonists are being considered as potential pharmacotherapeutics for OUD, the lack of cataleptic effects following D_3_R antagonism alone or in combination with morphine adds to accumulating evidence that D_3_R antagonists exhibit a desirable safety profile and lack adverse motoric effects as compared to nonselective D_2_-like receptor antagonists, even when opioids are concurrently administered.

In summary, we show that pretreatment with the selective D_3_R antagonist PG01037 attenuates acute morphine-induced hyperactivity similar to other selective D_3_R antagonists, indicating that reduction of the abuse-related effects of opioids may be a feature shared by all compounds in this drug class. Furthermore, the absence of cataleptic effects following administration of a D_3_R antagonist, alone or in combination with morphine, lends further support to their potential use and safety as treatments for OUD. Recently-developed, highly-selective D_3_R antagonists such as R-VK4-40 and VK4-116 exhibit the most desirable behavioral profiles for clinical investigation because they reduce the abuse-related behavioral effects of opioids and simultaneously do not disrupt their analgesic efficacy [32]. However, our present results with PG01037 provide a cautionary note that the potentiation of opioid-induced analgesia observed with R-VK4-40 and VK4-116 may not be universal for all D_3_R antagonists. Additional research will be needed to elucidate the neurobiological mechanisms by which highly-selective D_3_R antagonists such as R-VK4-40 and VK4-116 favorably alter the abuse-related and analgesic effects of opioids.

## ABBREVIATIONS

D_1_R: D_1_ receptor
D_2_R: D_2_ receptor
D_3_R: D_3_ receptor
D_4_R: D_4_ receptor
D_5_R: D_5_ receptor; receptor
DA: dopamine
NAc: nucleus accumbens
OUD: pioid use disorders
VTA: ventral tegmental area

## ACKNOWLEDGEMENTS

The authors thank Ms. J. Cao in the Medicinal Chemistry Section, NIDA-IRP for synthesis of PG01037.

## DATA STATEMENT

The datasets generated and analyzed in the current study are available from the corresponding author upon reasonable request.

## REFERENCES

[1] N.D. Volkow, C. Blanco, The changing opioid crisis: development, challenges and opportunities, Mol Psychiatry 26(1) (2021) 218–233.

[2] C. Blanco, N.D. Volkow, Management of opioid use disorder in the USA: present status and future directions, Lancet 393(10182) (2019) 1760–1772.

[3] M.J. Kreek, B. Reed, E.R. Butelman, Current status of opioid addiction treatment and related preclinical research, Sci Adv 5(10) (2019) eaax9140.

[4] G. Di Chiara, R.A. North, Neurobiology of opiate abuse, Trends Pharmacol Sci 13(5) (1992) 185–93.

[5] R.C. Pierce, V. Kumaresan, The mesolimbic dopamine system: the final common pathway for the reinforcing effect of drugs of abuse?, Neurosci Biobehav Rev 30(2) (2006) 215–38.

[6] R.A. Wise, Opiate reward: sites and substrates, Neurosci Biobehav Rev 13(2-3) (1989) 129–33.

[7] A. Bjorklund, S.B. Dunnett, Dopamine neuron systems in the brain: an update, Trends Neurosci 30(5) (2007) 194–202.

[8] R.Y. Moore, F.E. Bloom, Central catecholamine neuron systems: anatomy and physiology of the dopamine systems, Annu Rev Neurosci 1 (1978) 129–69.

[9] V.I. Chefer, B.L. Kieffer, T.S. Shippenberg, Basal and morphine-evoked dopaminergic neurotransmission in the nucleus accumbens of MOR- and DOR-knockout mice, Eur J Neurosci 18(7) (2003) 1915–22.

[10] K. Gysling, R.Y. Wang, Morphine-induced activation of A10 dopamine neurons in the rat, Brain Res 277(1) (1983) 119–27.

[11] D.P. Devine, P. Leone, D. Pocock, R.A. Wise, Differential involvement of ventral tegmental mu, delta and kappa opioid receptors in modulation of basal mesolimbic dopamine release: in vivo microdialysis studies, J Pharmacol Exp Ther 266(3) (1993) 1236–46.

[12] R. Spanagel, A. Herz, T.S. Shippenberg, Opposing tonically active endogenous opioid systems modulate the mesolimbic dopaminergic pathway, Proc Natl Acad Sci U S A 89(6) (1992) 2046–50.

[13] P. Leone, D. Pocock, R.A. Wise, Morphine-dopamine interaction: ventral tegmental morphine increases nucleus accumbens dopamine release, Pharmacol Biochem Behav 39(2) (1991) 469–72.

[14] S.W. Johnson, R.A. North, Opioids excite dopamine neurons by hyperpolarization of local interneurons, J Neurosci 12(2) (1992) 483–8.

[15] A. Matsui, J.T. Williams, Opioid-sensitive GABA inputs from rostromedial tegmental nucleus synapse onto midbrain dopamine neurons, J Neurosci 31(48) (2011) 17729–35.

[16] T.H. Hand, K.B. Franklin, 6-OHDA lesions of the ventral tegmental area block morphine-induced but not amphetamine-induced facilitation of self-stimulation, Brain Res 328(2) (1985) 233–41.

[17] A.E. Kelley, L. Stinus, S.D. Iversen, Interactions between D-ala-met-enkephalin, A10 dopaminergic neurones, and spontaneous behaviour in the rat, Behav Brain Res 1(1) (1980) 3–24.

[18] A.G. Phillips, F.G. LePiane, H.C. Fibiger, Dopaminergic mediation of reward produced by direct injection of enkephalin into the ventral tegmental area of the rat, Life Sci 33(25) (1983) 2505–11.

[19] J.E. Smith, G.F. Guerin, C. Co, T.S. Barr, J.D. Lane, Effects of 6-OHDA lesions of the central medial nucleus accumbens on rat intravenous morphine self-administration, Pharmacol Biochem Behav 23(5) (1985) 843–9.

[20] C. Spyraki, H.C. Fibiger, A.G. Phillips, Attenuation of heroin reward in rats by disruption of the mesolimbic dopamine system, Psychopharmacology (Berl) 79(2-3) (1983) 278–83.

[21] L. Stinus, G.F. Koob, N. Ling, F.E. Bloom, M. Le Moal, Locomotor activation induced by infusion of endorphins into the ventral tegmental area: evidence for opiate-dopamine interactions, Proc Natl Acad Sci U S A 77(4) (1980) 2323–7.

[22] B. Wang, F. Luo, X. Ge, A. Fu, J. Han, Effect of 6-OHDA lesions of the dopaminergic mesolimbic system on drug priming induced reinstatement of extinguished morphine CPP in rats, Beijing Da Xue Xue Bao Yi Xue Ban 35(5) (2003) 449–52.

[23] T.S. Shippenberg, R. Bals-Kubik, A. Herz, Examination of the neurochemical substrates mediating the motivational effects of opioids: role of the mesolimbic dopamine system and D-1 vs. D-2 dopamine receptors, J Pharmacol Exp Ther 265(1) (1993) 53–9.

[24] J.M. Beaulieu, R.R. Gainetdinov, The physiology, signaling, and pharmacology of dopamine receptors, Pharmacol Rev 63(1) (2011) 182–217.

[25] D.J. Reiner, I. Fredriksson, O.M. Lofaro, J.M. Bossert, Y. Shaham, Relapse to opioid seeking in rat models: behavior, pharmacology and circuits, Neuropsychopharmacology 44(3) (2019) 465–477.

[26] D.I. Cho, M. Zheng, K.M. Kim, Current perspectives on the selective regulation of dopamine D(2) and D(3) receptors, Arch Pharm Res 33(10) (2010) 1521–38.

[27] M.J. Millan, J.L. Peglion, J. Vian, J.M. Rivet, M. Brocco, A. Gobert, A. Newman-Tancredi, C. Dacquet, K. Bervoets, S. Girardon, et al., Functional correlates of dopamine D3 receptor activation in the rat in vivo and their modulation by the selective antagonist, (+)-S 14297: 1. Activation of postsynaptic D3 receptors mediates hypothermia, whereas blockade of D2 receptors elicits prolactin secretion and catalepsy, J Pharmacol Exp Ther 275(2) (1995) 885–98.

[28] M. Haney, A.S. Ward, R.W. Foltin, M.W. Fischman, Effects of ecopipam, a selective dopamine D1 antagonist, on smoked cocaine self-administration by humans, Psychopharmacology (Berl) 155(4) (2001) 330–7.

[29] T. Kishi, Y. Matsuda, N. Iwata, C.U. Correll, Antipsychotics for cocaine or psychostimulant dependence: systematic review and meta-analysis of randomized, placebo-controlled trials, J Clin Psychiatry 74(12) (2013) e1169–80.

[30] C.A. Heidbreder, A.H. Newman, Current perspectives on selective dopamine D(3) receptor antagonists as pharmacotherapeutics for addictions and related disorders, Ann N Y Acad Sci 1187 (2010) 4–34.

[31] P. Sokoloff, B. Le Foll, The dopamine D_3_ receptor, a quarter century later, Eur J Neurosci 45(1) (2017) 2–19.

[32] E. Galaj, A.H. Newman, Z.X. Xi, Dopamine D3 receptor-based medication development for the treatment of opioid use disorder: Rationale, progress, and challenges, Neurosci Biobehav Rev 114 (2020) 38–52.

[33] C.A. Boateng, O.M. Bakare, J. Zhan, A.K. Banala, C. Burzynski, E. Pommier, T.M. Keck, P. Donthamsetti, J.A. Javitch, R. Rais, B.S. Slusher, Z.X. Xi, A.H. Newman, High Affinity Dopamine D3 Receptor (D3R)-Selective Antagonists Attenuate Heroin Self-Administration in Wild-Type but not D3R Knockout Mice, J Med Chem 58(15) (2015) 6195–213.

[34] E. Galaj, M. Manuszak, S. Babic, S. Ananthan, R. Ranaldi, The selective dopamine D3 receptor antagonist, SR 21502, reduces cue-induced reinstatement of heroin seeking and heroin conditioned place preference in rats, Drug Alcohol Depend 156 (2015) 228–233.

[35] R. Hu, R. Song, R. Yang, R. Su, J. Li, The dopamine D(3) receptor antagonist YQA14 that inhibits the expression and drug-prime reactivation of morphine-induced conditioned place preference in rats, Eur J Pharmacol 720(1-3) (2013) 212–7.

[36] Y. Lv, R.R. Hu, M. Jing, T.Y. Zhao, N. Wu, R. Song, J. Li, G. Hu, Selective dopamine D3 receptor antagonist YQA14 inhibits morphine-induced behavioral sensitization in wild type, but not in dopamine D3 receptor knockout mice, Acta Pharmacol Sin 40(5) (2019) 583–588.

[37] Z.B. You, G.H. Bi, E. Galaj, V. Kumar, J. Cao, A. Gadiano, R. Rais, B.S. Slusher, E.L. Gardner, Z.X. Xi, A.H. Newman, Dopamine D3R antagonist VK4-116 attenuates oxycodone self-administration and reinstatement without compromising its antinociceptive effects, Neuropsychopharmacology 44(8) (2019) 1415–1424.

[38] C.J. Jordan, B. Humburg, M. Rice, G.H. Bi, Z.B. You, A.B. Shaik, J. Cao, A. Bonifazi, A. Gadiano, R. Rais, B. Slusher, A.H. Newman, Z.X. Xi, The highly selective dopamine D3R antagonist, R-VK4-40 attenuates oxycodone reward and augments analgesia in rodents, Neuropharmacology 158 (2019) 107597.

[39] G. de Guglielmo, M. Kallupi, S. Sedighim, A.H. Newman, O. George, Dopamine D3 Receptor Antagonism Reverses the Escalation of Oxycodone Self-administration and Decreases Withdrawal-Induced Hyperalgesia and Irritability-Like Behavior in Oxycodone-Dependent Heterogeneous Stock Rats, Front Behav Neurosci 13 (2019) 292.

[40] P. Grundt, E.E. Carlson, J. Cao, C.J. Bennett, E. McElveen, M. Taylor, R.R. Luedtke, A.H. Newman, Novel heterocyclic trans olefin analogues of N-{4-[4-(2,3-dichlorophenyl)piperazin-1-yl]butyl}arylcarboxamides as selective probes with high affinity for the dopamine D3 receptor, J Med Chem 48(3) (2005) 839–48.

[41] D.F. Manvich, A.K. Petko, R.C. Branco, S.L. Foster, K.A. Porter-Stransky, K.A. Stout, A.H. Newman, G.W. Miller, C.A. Paladini, D. Weinshenker, Selective D2 and D3 receptor antagonists oppositely modulate cocaine responses in mice via distinct postsynaptic mechanisms in nucleus accumbens, Neuropsychopharmacology 44(8) (2019) 1445–1455.

[42] C.J. Jordan, B.A. Humburg, E.B. Thorndike, A.B. Shaik, Z.X. Xi, M.H. Baumann, A.H. Newman, C.W. Schindler, Newly Developed Dopamine D3 Receptor Antagonists, R-VK4-40 and R-VK4-116, Do Not Potentiate Cardiovascular Effects of Cocaine or Oxycodone in Rats, J Pharmacol Exp Ther 371(3) (2019) 602–614.

[43] C.L. Broekkamp, A.G. Phillips, A.R. Cools, Stimulant effects of enkephalin microinjection into the dopaminergic A10 area, Nature 278(5704) (1979) 560–2.

[44] J.M. Delfs, L. Schreiber, A.E. Kelley, Microinjection of cocaine into the nucleus accumbens elicits locomotor activation in the rat, J Neurosci 10(1) (1990) 303–10.

[45] P.W. Kalivas, E. Widerlov, D. Stanley, G. Breese, A.J. Prange, Jr., Enkephalin action on the mesolimbic system: a dopamine-dependent and a dopamine-independent increase in locomotor activity, J Pharmacol Exp Ther 227(1) (1983) 229–37.

[46] P.H. Kelly, S.D. Iversen, Selective 6OHDA-induced destruction of mesolimbic dopamine neurons: abolition of psychostimulant-induced locomotor activity in rats, Eur J Pharmacol 40(1) (1976) 45–56.

[47] P.H. Kelly, P.W. Seviour, S.D. Iversen, Amphetamine and apomorphine responses in the rat following 6-OHDA lesions of the nucleus accumbens septi and corpus striatum, Brain Res 94(3) (1975) 507–22.

[48] V. Kumar, A. Bonifazi, M.P. Ellenberger, T.M. Keck, E. Pommier, R. Rais, B.S. Slusher, E. Gardner, Z.B. You, Z.X. Xi, A.H. Newman, Highly Selective Dopamine D3 Receptor (D3R) Antagonists and Partial Agonists Based on Eticlopride and the D3R Crystal Structure: New Leads for Opioid Dependence Treatment, J Med Chem 59(16) (2016) 7634–50.

[49] J.A. Kiritsy-Roy, S.M. Standish, L.C. Terry, Dopamine D-1 and D-2 receptor antagonists potentiate analgesic and motor effects of morphine, Pharmacol Biochem Behav 32(3) (1989) 717–21.

[50] M. Rodriguez-Arias, I. Broseta, M.A. Aguilar, J. Minarro, Lack of specific effects of selective D(1) and D(2) dopamine antagonists vs. risperidone on morphine-induced hyperactivity, Pharmacol Biochem Behav 66(1) (2000) 189–97.

[51] R.E. Wilcox, M. Bozarth, R.A. Levitt, Reversal of morphine-induced catalepsy by naloxone microinjections into brain regions with high opiate receptor binding: a preliminary report, Pharmacol Biochem Behav 18(1) (1983) 51–4.

[52] P.R. Sanberg, M.D. Bunsey, M. Giordano, A.B. Norman, The catalepsy test: its ups and downs, Behav Neurosci 102(5) (1988) 748–59.

[53] A. Fink-Jensen, L.S. Schmidt, D. Dencker, C. Schulein, J. Wess, G. Wortwein, D.P. Woldbye, Antipsychotic-induced catalepsy is attenuated in mice lacking the M4 muscarinic acetylcholine receptor, Eur J Pharmacol 656(1-3) (2011) 39–44.

[54] Z.B. You, J.T. Gao, G.H. Bi, Y. He, C. Boateng, J. Cao, E.L. Gardner, A.H. Newman, Z.X. Xi, The novel dopamine D3 receptor antagonists/partial agonists CAB2-015 and BAK4-54 inhibit oxycodone-taking and oxycodone-seeking behavior in rats, Neuropharmacology 126 (2017) 190–199.

[55] L.M. Pritchard, A.H. Newman, R.K. McNamara, A.D. Logue, B. Taylor, J.A. Welge, M. Xu, J. Zhang, N.M. Richtand, The dopamine D3 receptor antagonist NGB 2904 increases spontaneous and amphetamine-stimulated locomotion, Pharmacol Biochem Behav 86(4) (2007) 718–26.

[56] E. Galaj, S. Ananthan, M. Saliba, R. Ranaldi, The effects of the novel DA D3 receptor antagonist SR 21502 on cocaine reward, cocaine seeking and cocaine-induced locomotor activity in rats, Psychopharmacology (Berl) 231(3) (2014) 501–10.

[57] R. Song, R.F. Yang, N. Wu, R.B. Su, J. Li, X.Q. Peng, X. Li, J. Gaal, Z.X. Xi, E.L. Gardner, YQA14: a novel dopamine D3 receptor antagonist that inhibits cocaine self-administration in rats and mice, but not in D3 receptor-knockout mice, Addict Biol 17(2) (2012) 259–73.

[58] S. Schildein, A. Agmo, J.P. Huston, R.K. Schwarting, Intraaccumbens injections of substance P, morphine and amphetamine: effects on conditioned place preference and behavioral activity, Brain Res 790(1-2) (1998) 185–94.

[59] F.J. Vaccarino, W.A. Corrigall, Effects of opiate antagonist treatment into either the periaqueductal grey or nucleus accumbens on heroin-induced locomotor activation, Brain Res Bull 19(5) (1987) 545–9.

[60] H. Teitelbaum, P. Giammatteo, G.A. Mickley, Differential effects of localized lesions of n. accumbens on morphine- and amphetamine-induced locomotor hyperactivity in the C57BL/6J mouse, J Comp Physiol Psychol 93(4) (1979) 745–51.

[61] P.W. Kalivas, P. Duffy, Sensitization to repeated morphine injection in the rat: possible involvement of A10 dopamine neurons, J Pharmacol Exp Ther 241(1) (1987) 204–12.

[62] L.J. Vanderschuren, P.W. Kalivas, Alterations in dopaminergic and glutamatergic transmission in the induction and expression of behavioral sensitization: a critical review of preclinical studies, Psychopharmacology (Berl) 151(2-3) (2000) 99–120.

[63] P. Vezina, P.W. Kalivas, J. Stewart, Sensitization occurs to the locomotor effects of morphine and the specific mu opioid receptor agonist, DAGO, administered repeatedly to the ventral tegmental area but not to the nucleus accumbens, Brain Res 417(1) (1987) 51–8.

[64] K. Hattori, S. Uchino, T. Isosaka, M. Maekawa, M. Iyo, T. Sato, S. Kohsaka, T. Yagi, S. Yuasa, Fyn is required for haloperidol-induced catalepsy in mice, J Biol Chem 281(11) (2006) 7129–35.

[65] M.J. Millan, A. Dekeyne, J.M. Rivet, T. Dubuffet, G. Lavielle, M. Brocco, S33084, a novel, potent, selective, and competitive antagonist at dopamine D(3)-receptors: II. Functional and behavioral profile compared with GR218,231 and L741,626, J Pharmacol Exp Ther 293(3) (2000) 1063–73.

[66] C. Achat-Mendes, P. Grundt, J. Cao, D.M. Platt, A.H. Newman, R.D. Spealman, Dopamine D3 and D2 receptor mechanisms in the abuse-related behavioral effects of cocaine: studies with preferential antagonists in squirrel monkeys, J Pharmacol Exp Ther 334(2) (2010) 556–65.

[67] C. Reavill, S.G. Taylor, M.D. Wood, T. Ashmeade, N.E. Austin, K.Y. Avenell, I. Boyfield, C.L. Branch, J. Cilia, M.C. Coldwell, M.S. Hadley, A.J. Hunter, P. Jeffrey, F. Jewitt, C.N. Johnson, D.N. Jones, A.D. Medhurst, D.N. Middlemiss, D.J. Nash, G.J. Riley, C. Routledge, G. Stemp, K.M. Thewlis, B. Trail, A.K. Vong, J.J. Hagan, Pharmacological actions of a novel, high-affinity, and selective human dopamine D(3) receptor antagonist, SB-277011-A, J Pharmacol Exp Ther 294(3) (2000) 1154–65.

