## Supplemental Figures 1 and 2 for "Effects of the selective dopamine D_3_ receptor antagonist PG01037 on morphine-induced hyperactivity and antinociception in mice"

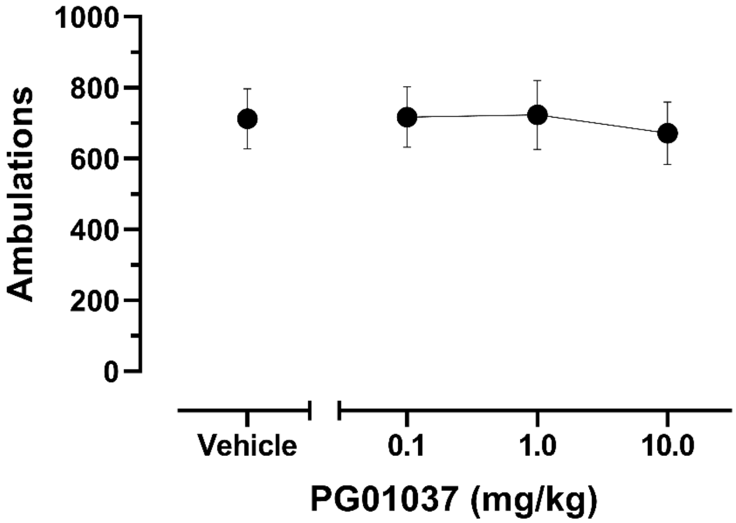


**Fig. S1** Effects of PG01037 on basal locomotor activity in the 30-min period prior to morphine administration. Mice were pretreated with vehicle or 0.1 – 10 mg/kg PG01037, followed 30 min later by vehicle or 5.6 – 56 mg/kg morphine. Shown are mean ± SEM total ambulations in the 30-min period following PG01037 administration, prior to morphine injection. Because mice were administered each dose of PG01037 on three separate occasions (i.e. prior to three different doses of morphine), ambulations were averaged across the three administrations for each dose of PG01037 for each subject. All mice received all treatments (n = 16). Doses on the abscissa are plotted along a log scale.

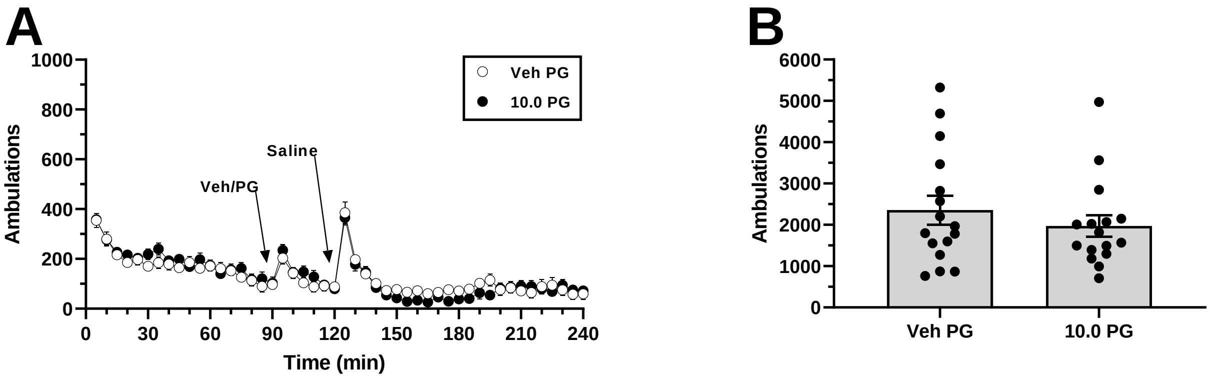


**Fig. S2** Basal locomotor activity is not altered by PG01037 in a 150-min duration following its administration. **A** Time course. PG01037 (vehicle, 10 mg/kg) was administered 30 min prior to saline. Shown are mean ± SEM ambulations recorded in 5-min bins. Arrows indicate time of pretreatment and saline injections. **B** Shown are total ambulations in the 120-min period following saline injection. Gray bars represent mean ± SEM total ambulations, with values from individual subjects shown in the superimposed scatterplot. All mice received both treatments (n = 16). “Veh” = vehicle; “PG” = PG01037.
